# Harmonizing Functional Connectivity Reduces Scanner Effects in Community Detection

**DOI:** 10.1101/2021.12.03.469269

**Authors:** Andrew A. Chen, Dhivya Srinivasan, Raymond Pomponio, Yong Fan, Ilya M. Nasrallah, Susan M. Resnick, Lori L. Beason-Held, Christos Davatzikos, Theodore D. Satterthwaite, Dani S. Bassett, Russell T. Shinohara, Haochang Shou

## Abstract

Community detection on graphs constructed from functional magnetic resonance imaging (fMRI) data has led to important insights into brain functional organization. Large studies of brain community structure often include images acquired on multiple scanners across different studies. Differences in scanner can introduce variability into the downstream results, and these differences are often referred to as scanner effects. Such effects have been previously shown to significantly impact common network metrics. In this study, we identify scanner effects in data-driven community detection results and related network metrics. We assess a commonly employed harmonization method and propose new methodology for harmonizing functional connectivity that leverage existing knowledge about network structure as well as patterns of covariance in the data. Finally, we demonstrate that our new methods reduce scanner effects in community structure and network metrics. Our results highlight scanner effects in studies of brain functional organization and provide additional tools to address these unwanted effects. These findings and methods can be incorporated into future functional connectivity studies, potentially preventing spurious findings and improving reliability of results.

## 1 Introduction

To better understand the functional organization of the brain, researchers have used functional imaging to observe task-associated and resting-state brain activity at high temporal resolutions. These measurements can be used to construct brain networks, which describe groups of regions that are strongly linked and are jointly associated with unique neurophysiological functions (Salvador et al., 2005). Community detection methods provide a data-driven approach to examine these brain networks by identifying groups of tightly coupled regions called communities. Abnormalities in specific brain communities have been associated with Alzheimer’s disease and major depressive disorder among other illnesses (Dragomir et al., 2019; He et al., 2018). Owing to their contributions to significant clinical findings, functional imaging and community detection metrics have been considered for their suitability as potential biomarkers for neurological and psychiatric disorders (Gallen & D’Esposito, 2019; Hohenfeld et al., 2018; Parkes et al., 2020). Understanding confounding factors that can impact community detection results is thus a particularly urgent task.

Functional imaging studies have sought larger sample sizes by collecting data across multiple institutions and scanners, which potentially introduces variation due to scanner referred to as scanner effects. Several studies have characterized and attempted to address these effects, but do not examine how scanner effects can influence brain network organization and relevant metrics (Dansereau et al., 2017; Feis et al., 2015; Friedman et al., 2006). To the best of our knowledge, only one study has investigated the influence of scanner effects on brain network constructed using resting-state functional magnetic resonance imaging (rsfMRI) data and proposed a method for harmonization by adapting ComBat (Yu et al., 2018), a method initially proposed in gene expression studies for batch effect correction and adapted in neuroimaging to effectively mitigate scanner effects (Johnson et al., 2007; Fortin et al., 2017, 2018). To our knowledge, no study has investigated how community detection could be influenced by differences in scanners.

In this study, we assess whether scanner effects can drive differences in estimated communities using data from two large-scale multi-site rsfMRI studies. We first determine that scanner can strongly influence community detection results in both population-representative networks and subject-specific networks. We then examine community metrics and identify notable scanner effects. To address these issues, we propose new methodology and compare these two methods with the existing rsfMRI harmonization method and find that all methods reduce scanner effects, but our proposed methods provide the best results in several evaluations. Our results highlight the need to consider scanner effects in rsfMRI studies of brain functional organization and provide insight into mitigating these issues.

## 2 Methods

### 2.1 Image acquisition and processing

Our sample consists of fMRI scans from the iSTAGING consortium (Habes et al., 2021), which includes data from eleven different studies. For this study, we examine a subset of fMRI data from the Baltimore Longitudinal Study of Aging (BLSA; Shock, 1984) and the Coronary Artery Risk Development in Young Adults (CARDIA; Friedman et al., 1988) study. Through its extensive multimodal analysis, the BLSA study has provided marked insight into the physical and cognitive effects of aging. The fMRI data from BLSA are particularly suited to our analysis since all images were acquired on a single scanner (see Table 1), allowing for direct comparison with CARDIA scanners. The CARDIA study aims to identify risk factors for cardiovascular disease and collected rsfMRI scans as part of its assessment of neurological outcomes. The CARDIA study has four scanners, but one scanner is excluded from our analysis since only four subjects are acquired on it.

**Table 1:** Demographics by scanner for the four scanners across the two included studies. The study sample consists of subjects from the Baltimore Longitudinal Study of Aging (BLSA) and the Coronary Artery Risk Development in Young Adults (CARDIA) studies. We include the one scanner from the BLSA study and three scanners from the CARDIA study in our analyses. ANOVA *p*-values for testing differences in the mean of continuous variables and Chi-squared test *p*-values for testing the differences in categorical variables are reported in the rightmost column.

|  | BLSA | CARDIA 1 | CARDIA 3 | CARDIA 4 | p |
| --- | --- | --- | --- | --- | --- |
| Scanner | Philips Achieva 3T | Philips Achieva 3T | Siemens Trio 3T | Siemens Trio 3T |  |
| Number of Subjects | 357 | 57 | 185 | 139 |  |
| Age (mean (SD)) | 65.42 (15.25) | 50.12 (3.42) | 50.30 (3.31) | 50.12 (3.64) | <0.01 |
| Male (%) | 161 (45.1) | 22 (38.6) | 79 (42.7) | 55 (39.6) | 0.62 |

The resting state fMRI images for BLSA are acquired using Philips Achieva 3T scanner with repetition time (TR) of 2 seconds and echo time (TE) of 30 ms, with a total duration per scan of 6 minutes. Other parameters include a field of view (FOV) of 80 × 80mm, 37 axial slices, 3 × 3mm voxel resolution, and 180 time frames. CARDIA scans are acquired across four different sites using similar protocols, with a TR of 2 seconds, TE of 25 ms, FOV of 224 × 224 mm, 64 axial slices, slice thickness of 3.5 mm, and 120 time frames. All the functional images are slice time-corrected. MCFLIRT (Jenkinson et al., 2002) is used to realign all volumes the selected reference volume. Time series are band-pass filtered to retain frequencies between 0.01 – 0.08 Hz. All pre-processing steps were carried out using FSL FEAT (www.fmrib.ox.ac.uk/fsl).

Besides band-pass filtering and usual motion correction, mean relative displacement, relative voxel-wise motion displacement and DVARS (Derivative of rms VARiance over voxelS) are calculated to summarize whole brain signal change. We apply a confound regression procedure with scrubbing (Power et al., 2012) using a 36-parameter model including motion, white matter, cerebrospinal fluid (CSF), global signal time courses, and temporal derivatives of these regressors. Subjects whose mean relative displacement greater than 0.2 mm are flagged and excluded. Filtered cleaned time series are then co-registered with structural MRI using DRAMMS (Deformable Registration via Attribute Matching and Mutual-Saliency Weighting; Ou et al., 2011) affine-only and then transformed to standard MNI space with 2× 2× 2mm resolution by combining warps from structural to Montreal Neurological Institute (MNI) space.

Following all pre-processing steps, functional connectivity matrices of dimension 264 × 264 were obtained by calculating Pearson correlations between the filtered, cleaned time courses extracted from each pair of ROIs defined by the Power atlas (Power et al., 2011). These 264 ROIs are grouped into 14 distinct subnetworks based on communities derived using Infomap (Power et al., 2011).

### 2.2 Functional connectivity ComBat

Across gene expression data and several imaging modalities, ComBat has proven to be an effective harmonization method which uses empirical Bayes to leverage batch effect information across features (Johnson et al., 2007; Fortin et al., 2017, 2018). ComBat has been applied to functional connectivity (FC) and shown to substantially reduce scanner effects in several network metrics including global efficiency, local efficiency, and within-community mean edge weights (Yu et al., 2018). Following a standard network model (Bassett et al., 2018), we first compute connectivity matrices from ROI-level time series using Pearson correlation between each time series. The correlation values are Fisher-transformed to range across all real numbers, which are subsequently used as edge weights in constructing networks from the connectivity matrices (Bassett et al., 2018). Let ***y**_ij_* = (*y*_*ij*1_, *y*_*ij*2_,…, *y_ijv_*)*^T^*, *i* = 1, 2,…, *K*, *j* = 1, 2,…, *n_i_* denote the *v*-dimensional Fisher-transformed edge weights where *i* indexes scanner, *j* indexes subjects within scanners, *n_i_* is the number of subjects acquired on scanner *i*, and *V* is the number of features. We aim to harmonize the data from these 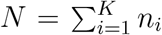 subjects across the *K* scanners. FC-ComBat (Yu et al., 2018) assumes that the *V* edges *v* = 1, 2,…, *V* follow

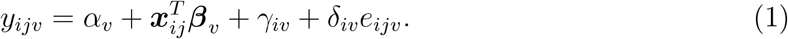

Using least-squares estimates for the regression coefficients, we first standardize the data as

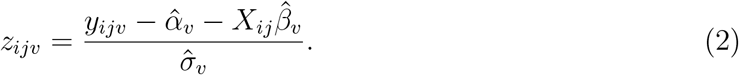

Then, we obtain empirical Bayes estimates for the scanner parameters, to yield harmonized edges as

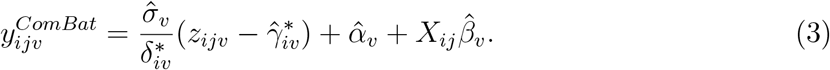

These harmonized edge weights can be used to construct adjacency matrices for downstream network analyses.

### 2.3 Proposed methodologies

FC-ComBat assumes that scanner effects influence all edges similarly and uses empirical Bayes to borrow information across all edges. However, prior network analyses may indicate distinct scanner effects within-network and between-network edges. By assuming scanner effects are shared across all edges, FC-ComBat may not adequately reduce network-specific scanner differences.

We propose and compare two novel methods that harmonizes functional connectivity data by utilizing correlations within and between networks. The first method is Block-ComBat (Bl-ComBat), which uses information across within-network edges by specifying subnetworks prior to harmonization. Our method first applies FC-ComBat followed by additional ComBat steps within each block. Our harmonization selectively borrows within-network information to address scanner effects that may not feature prominently in the first ComBat step. These blocks can be derived from prior knowledge of brain subnetworks, often obtained through structural and functional brain parcellation or clustering.

Community detection algorithms leverage patterns among groups of edges, which tend to be stronger within the same community and weaker between distinct communities (Newman, 2006). Scanner effects in groups of edges could thus impact community detection algorithms. ComBat harmonizes the mean and variance of each edge separately, and does not directly address effects in a group of edges. However, our recent harmonization method called CovBat addresses scanner effects in covariance by simultaneously harmonizing groups of features in principal component directions and may be well-suited to addressing these scanner effects (Chen et al., 2021).

Our second method is functional connectivity CovBat (FC-CovBat), which utilizes Cov-Bat as the primary harmonization step. Our method uses the same Fisher-transformation and vectorization of edges as in FC-ComBat. We again assume that the Fisher-transformed edges follow

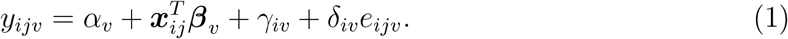

However, the error vectors ***e**_ij_* = (*e*_*ij*1_, *e*_*ij*2_,…, *e_ijp_*)*^T^* ~ *N*(**0**, Σ*_i_*) may be spatially correlated and differ in covariance across scanner. We first perform ComBat to remove the mean and variance shifts in the marginal distributions of the cortical thickness measures. Then, we additionally residualize with respect to the intercept and covariates to obtain ComBat-adjusted residuals denoted 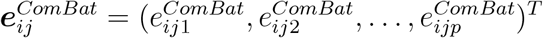 where *p* is the number of features. We define these residuals using previous notation as

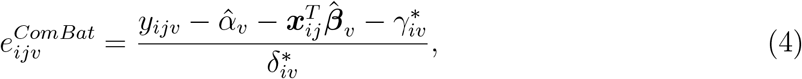

where *i* = 1, 2,…, *M*, *j* = 1,2,…, *n_i_*, *M* is the number of scanners, and *n_i_* is the number of subjects acquired at scanner *i*, and the variables 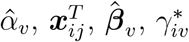, and 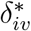 are the same as in FC-ComBat.

The 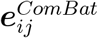 are assumed to have mean 0 and their covariance matrices which we denote by Σ*_i_* may differ across sites. CovBat performs principal components analysis (PCA) on the full data residuals and represents the full data covariance matrix as 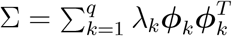 where the rank 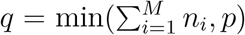, *λ_k_* are the eigenvalues of Σ, and ***ϕ**_k_* are the principal compo-nents obtained as the eigenvectors of Σ. PCA is performed on the sample covariance matrix 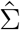 to obtain estimated eigenvalues 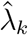 and eigenvectors 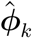. The ComBat-adjusted residuals can then be expressed as 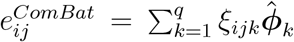 where *ξ_ijk_* are the principal component scores.

We approximate the within-site covariance matrices as 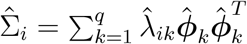 where 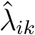 are within-site eigenvalues estimated as the sample variance of the principal component scores 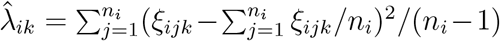. The 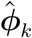 are estimated from the full data covariance. Then, we model scanner effects in the principal component scores as

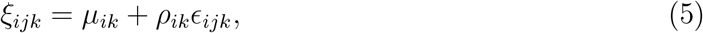

where 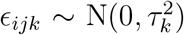, *τ_k_* is the standard deviation of the error, and *μ_ik_*, *ρ_ik_* are the center and scale parameters corresponding to principal components *k* = 1, 2,… *K* where *K* ≤ *q* is a tuning parameter chosen to capture the desired proportion of the variation in the observations. If *K* is chosen such that *K* = *q*, then all principal components are harmonized. We can then estimate each of the *K* pairs of center and scale parameters by finding the values that bring each site’s mean and variance in scores to the pooled mean and variance, which we denote 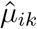) and 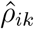. We then remove the site effect in the scores by subtracting out the estimated center parameter and dividing out the estimated scale parameter via 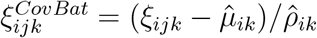.

We obtain CovBat-adjusted residuals 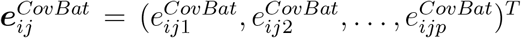 by projecting the adjusted scores back into the residual space via

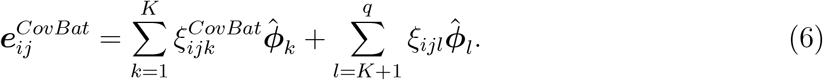

We then add the intercepts and covariates effects to obtain CovBat-adjusted edges

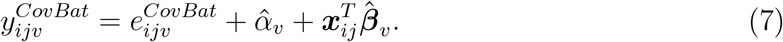

The CovBat methodology requires selection of the number of principal components to harmonize. Based on previous investigations, we choose to include principal components that explain 95% of the variation (Chen et al., 2021).

### 2.4 Evaluation of functional connectivity harmonization

Previous investigations on FC-ComBat examine the impact of harmonization on network metrics including mean edge weight within subnetworks, efficiency, and nodal strength (Yu et al., 2018). We extend these evaluations and propose four analyses that inform how harmonization can influence scanner effects in community detection. First, we regress edge weights and mean edge weights within and between subnetworks to determine how scanner effects influence the construction of networks. Second, we average functional connectivities acquired on the same scanner to determine how scanner effects influence communities derived from these population-average networks. Third, we find communities for each subject and use distance-based regression to assess scanner effects in subject-specific communities. Fourth, we compute network metrics related to community structure and examine associations with scanner and clinical variables. All four analyses are performed before and after harmonization using FC-ComBat, Bl-ComBat, and FC-CovBat to compare these three harmonization methods on their removal of scanner effects.

#### 2.4.1 Scanner effects in edge weights

Following previous investigations on FC-ComBat, we inspect the edge weights before and after harmonization to assess general changes in the raw values, removal of scanner effects, and preservation of biological variability. Scanner effects in these edge weights are likely to impact the downstream weighted networks. We first inspect the 34716-dimensional edge vectors using t-SNE (van der Maaten & Hinton, 2008) with a perplexity value of 30, which provides a low-dimensional visualization of the data that captures dominant patterns of variability. We then compare t-SNE plots before and after harmonization to visualize changes in the functional connectivity matrices.

To further assess the extent of scanner effects in the connectivity values, we perform linear regression for scanner and biological covariates on two scales and compare results before and after harmonization. First, we regress each edge separately on scanner, age, and sex to determine if scanner effects exist in individual edges. Second, we average within atlas subnetworks to obtain 14 within-network mean edge weight values and 91 between-network mean edge weights. Then, we perform regression on these mean edge weights, which are often used as outcomes in studies of functional connectivity (Yu et al., 2018; Varangis et al., 2019). We assess statistical significance controlling the false discovery rate at 5% (Benjamini & Hochberg, 1995).

#### 2.4.2 Population-average community detection

We next compare population-average community structure within each scanner to determine overall differences in community structure, similar to several studies contrasting communities between patient groups (Dragomir et al., 2019; He et al., 2018; Varangis et al., 2019). Our analysis of community structure utilizes two prominent algorithms that leverage both positive and negative edge weights. Following previous studies (Alexander-Bloch et al., 2012; Betzel et al., 2013; Varangis et al., 2019), we first average functional connectivities within each scanner then employ a version of the Louvain method (Blondel et al., 2008) adapted for signed networks using an asymmetric measure of modularity with positive and negative weights (Rubinov & Sporns, 2011), implemented in the Brain Connectivity Toolbox (BCT version 2019-03-03; Rubinov & Sporns (2010)). We repeat the algorithm 100 times and derive a consensus partition by applying the Louvain method to a consensus matrix constructed across partitions from the 100 runs, which reduces variability introduced by the algorithm (Lancichinetti & Fortunato, 2012). For all signed Louvain runs, we choose a tuning parameter *λ* to yield median 14 communities per scanner to approximate the number of communities in the atlas (Power et al., 2011). To evaluate community structure differences before and after harmonization, we inspect these scanner-average communities visually. For a quantitative evaluation, we also compute the adjusted Rand index (ARI; Hubert & Arabie, 1985)) for all pairwise comparisons between scanners and average within harmonization method to yield a mean ARI value that is lower if communities are more similar among scanners.

To assess whether scanner effects are present across multiple community detection methods, we also obtain communities using a weighted stochastic block model (WSBM; Aicher et al., 2013), which has found notable results applied to brain functional networks (Betzel et al., 2018). Since our networks are fully-connected, we assume that all edges exist and that the that edge weights are normally distributed with parameters varying across different pairs of nodal communities. We fix the number of communities at *K* =14 for comparison with the atlas and implement the WSBM using code available online (aaronclauset.github.io/wsbm; version 1.3). Similar to our signed Louvain evaluation, we compare community partitions across scanners visually and by examining the mean ARI among pairwise comparisons.

#### 2.4.3 Subject-specific communities

Examining communities determined via the average connectivity matrix within each scanner does not capture subject-to-subject variability in community structure, which may be influenced by scanner. To assess the existence of scanner effects on a subject level and determine if harmonization is adequate to address these, we use the signed Louvain algorithm to obtain subject-specific communities. First, we construct signed subject-level networks using the Fisher-transformed edge weights. For each subject, we then obtain a consensus partition across 100 runs of the signed Louvain algorithm, choosing a tuning parameter *λ* to yield a median of 14 communities across subjects.

To detect associations of these community partitions with scanner and biological variables, we use distance-based approaches that enable analysis of variance (ANOVA) and regression with an appropriate choice of distance measure. Previous studies have taken a similar approach by performing an ANOVA-like analysis with a similarity measure (Alexander-Bloch et al., 2012). We instead compute variation of information (VI) for each pair of subjects, which is a proper distance measure on the space of clusterings (Meilă, 2007). Sub-sequently, we perform permutational analysis of variance (PERMANOVA) to test for group differences and multivariate distance matrix regression (MDMR) to test for associations with covariates. We first apply PERMANOVA twice to first determine if communities differ across all scanner groups and then assess if differences exist among CARDIA scanners. The two studies included have considerably different age distributions (Table 1), which may confound our detection of scanner effects. To control for age and sex while evaluating associations with scanner, we apply MDMR while including scanner, age, and sex in the model.

#### 2.4.4 Computation of network metrics

ComBat has previously been shown to remove scanner effects in common network metrics including default mode network connectivity and efficiency values while preserving biological signals (Yu et al., 2018). We examine additional metrics that describe the community structure of signed networks. All metrics are calculated using the Brain Connectivity Toolbox (BCT) version 2019-03-03 (Rubinov & Sporns, 2010).

Modularity captures the extent to which a network segregates into a given community partition (Newman, 2006). Our experiment involves both communities derived from their brain atlas and from the signed Louvain algorithm. We compute modularity for both partitions and use linear regression to assess whether the Power atlas modularity (A. Mod.) and signed Louvain-derived community partition modularity (C. Mod.) are associated with scanner or age. Scanner effects in either case would demonstrate that the goodness-of-fit for relevant partitions could be confounded by scanner differences.

Clustering coefficient (CC) similarly captures the tendency for a network to form communities, but provides distinct information from modularity. CC operates on triplets of nodes and does not rely on a community partition, instead measuring local density in connectivity (Watts & Strogatz, 1998). Generalizations of the clustering coefficient have been developed for weighted (Onnela et al., 2005) and signed networks (Costantini & Perugini, 2014). We compute the signed clustering coefficient for each subject’s brain network and regress on scanner and age before and after harmonization.

A network metric that captures interrelatedness among nodes with respect to community assignments is the participation coefficient, which measures the relative strength of connections within network compared to outside the network (Guimerà & Amaral, 2005). A previous study revealed that the average participation coefficient can be influenced by age (Varangis et al., 2019). Following previous work (Zamani Esfahlani et al., 2020), we compute participation coefficient separately for positive edges (positive PC) and negative edges (negative PC). As previously noted, positive PC has a more direct interpretation as the relative strength of positive correlations with subnetworks (Zamani Esfahlani et al., 2020), so we use positive PC as the primary outcome. For each subnetwork, we summarize the positive and negative PCs by averaging across nodes within the subnetwork. As with other metrics, we assess scanner and age effects using linear regression.

## 3 Results

### 3.1 Subject characteristics

Our final dataset includes imaging data from 357 BLSA participants and 381 CARDIA participants. Table 1 shows the demographics by scanner, showing that BLSA and CARDIA 1 both are Philips Achieva 3T scanners while CARDIA 3 and 4 are Siemens Trio 3T scanners. There is a significant difference in age among participants across scanners (analysis of variance, *p* < 0.01), which is driven by the considerably older BLSA subjects with fMRI data. No significant differences are found in the sex proportion across scanners.

### 3.2 Functional connectivity associated with scanner

Inspection of the functional connectivity via t-SNE indicates that there is considerable separation across studies, with CARDIA observations staying roughly grouped together (Fig. 1). However, these visual differences could be the driven by differences in the age distribution (see Table 1 for details). We isolate associations with scanner by controlling for age and sex in our edge-wise regression and show scanner effects across a large portion of the within-network and between-network edges (Fig. 2). Fig. 3 displays results for within-network and between-network mean edge weights, further highlighting significant associations with scanner.

**Figure 1:**
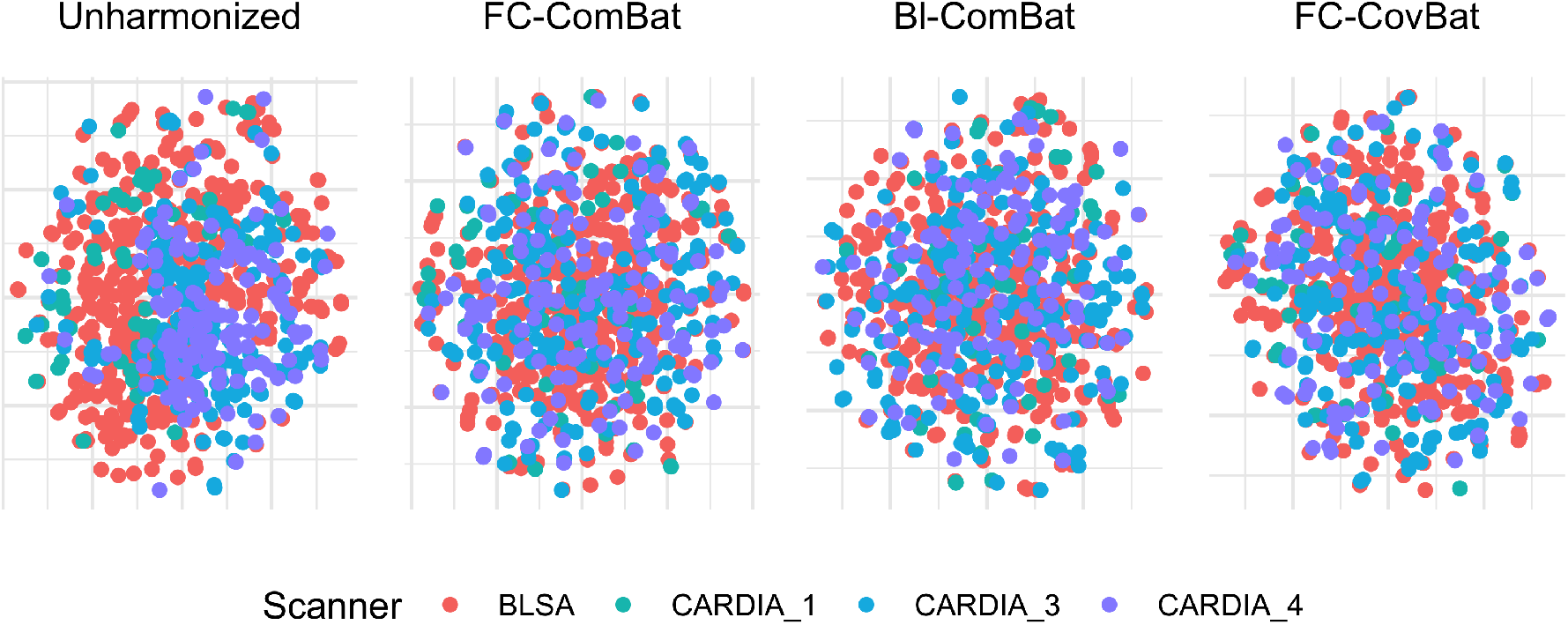
t-SNE visualization of functional connectivity matrices before and after harmonization. t-SNE is used to show segregation of unharmonized functional connectivity matrices into scanner-specific clusters. The matrices are shown to be more integrated after harmonization. t-SNE is performed using Frobenius distances between functional connectivity matrices as inputs. The perplexity value was chosen at 30.

**Figure 2:**
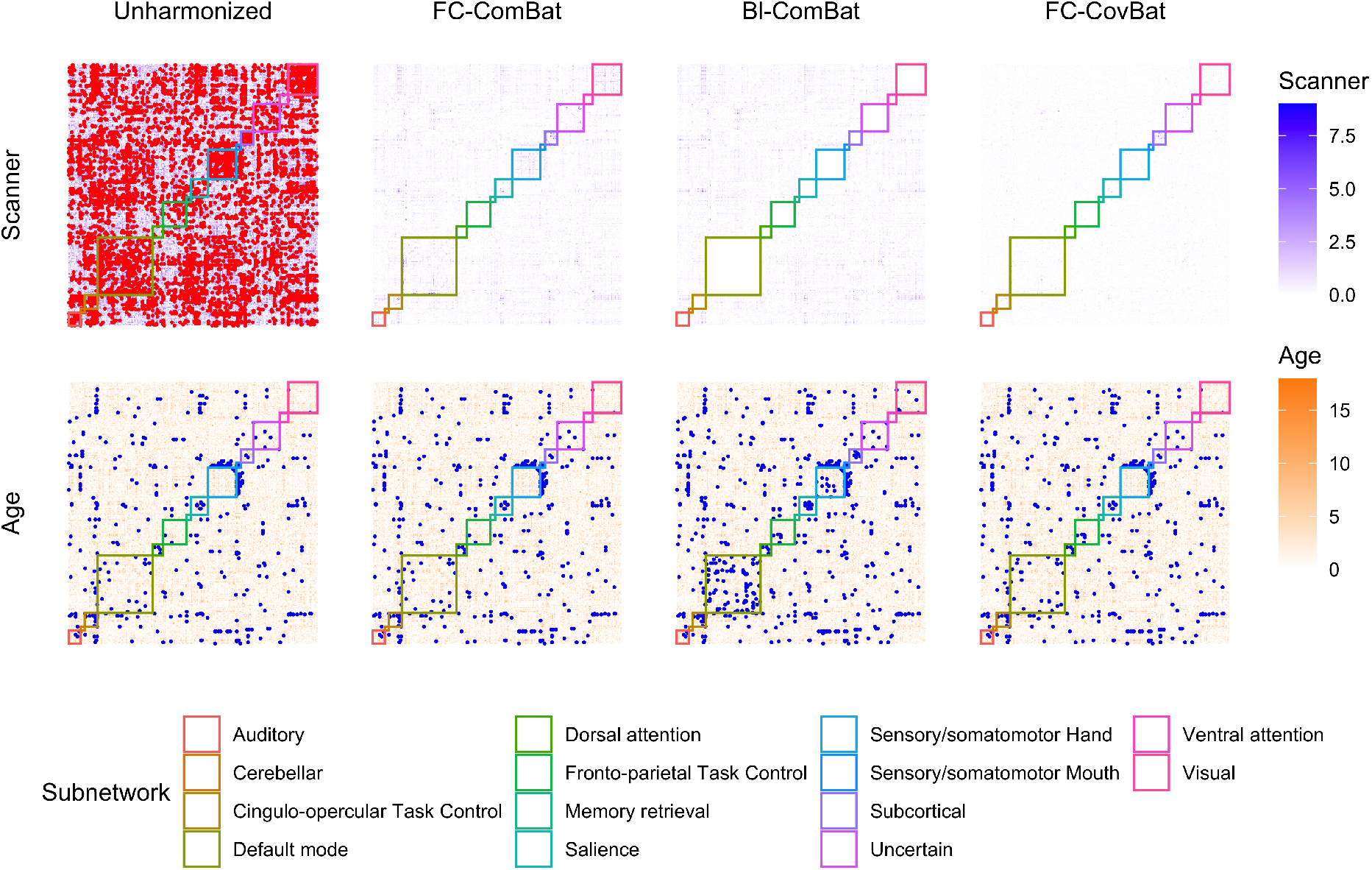
Edge-wise regression of functional connectivity before and after harmonization. Effects of harmonization on associations of edge weights with scanner and age are shown. Linear models are fit to each edge including scanner, age, and sex as covariates. Negative log *p*-values are shown where red and blue highlighting indicate significant edges for scanner and age respectively after false discovery rate control at the 0.05 level.

**Figure 3:**
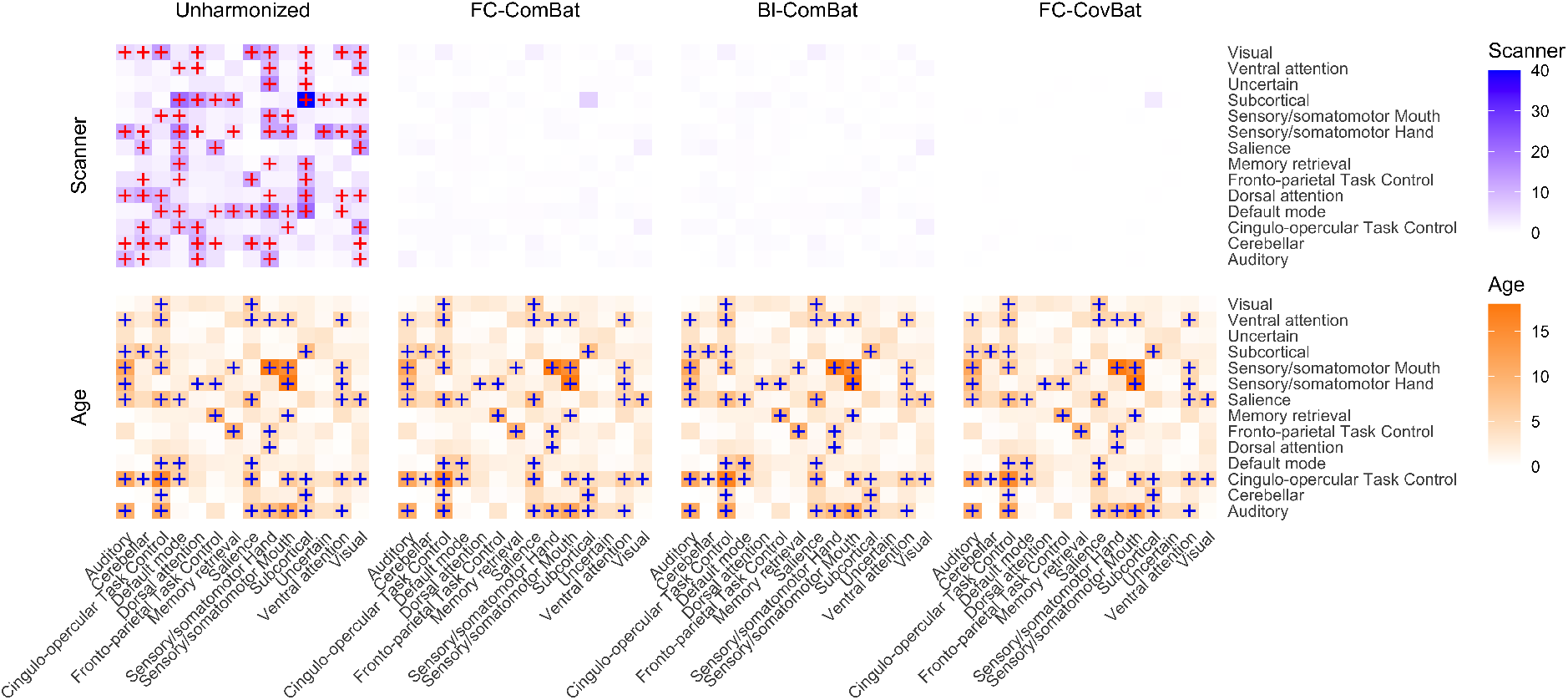
Regression of within-network and between-network mean edge weights. Significant associations of mean edge weights with scanner and age are shown before and after harmonization. Linear models are fit to each within and between connectivity value while including scanner, age, and sex as covariates. Negative log *p*-values are shown where red and blue highlighting indicate significant associations with scanner and age after false discovery rate control at the 0.05 level.

The tested harmonization methods all reduce separation of functional connectivity matrices across studies visualized through t-SNE in Fig. 1. The harmonization methods all remove scanner associations in both edges and mean edge weights while retaining associations with age (Fig. 2 and Fig. 3). Among the harmonization methods, Fig. 4 shows that Bl-ComBat provides the greatest reduction of the log-*p* values for within-network mean edge weights (Wilcoxon rank-sum test against FC-ComBat, *p* < 0.001; Wilcoxon rank-sum test against FC-CovBat *p* < 0.001) while FC-CovBat shows the best performance for between-network mean edge weights (Wilcoxon rank-sum test against FC-ComBat, *p* < 0.001; Wilcoxon ranksum test against Bl-ComBat *p* < 0.001). For age associations, FC-ComBat and FC-CovBat both retain unharmonized associations while Bl-ComBat recovers additional age associations for some within-network edges.

**Figure 4:**
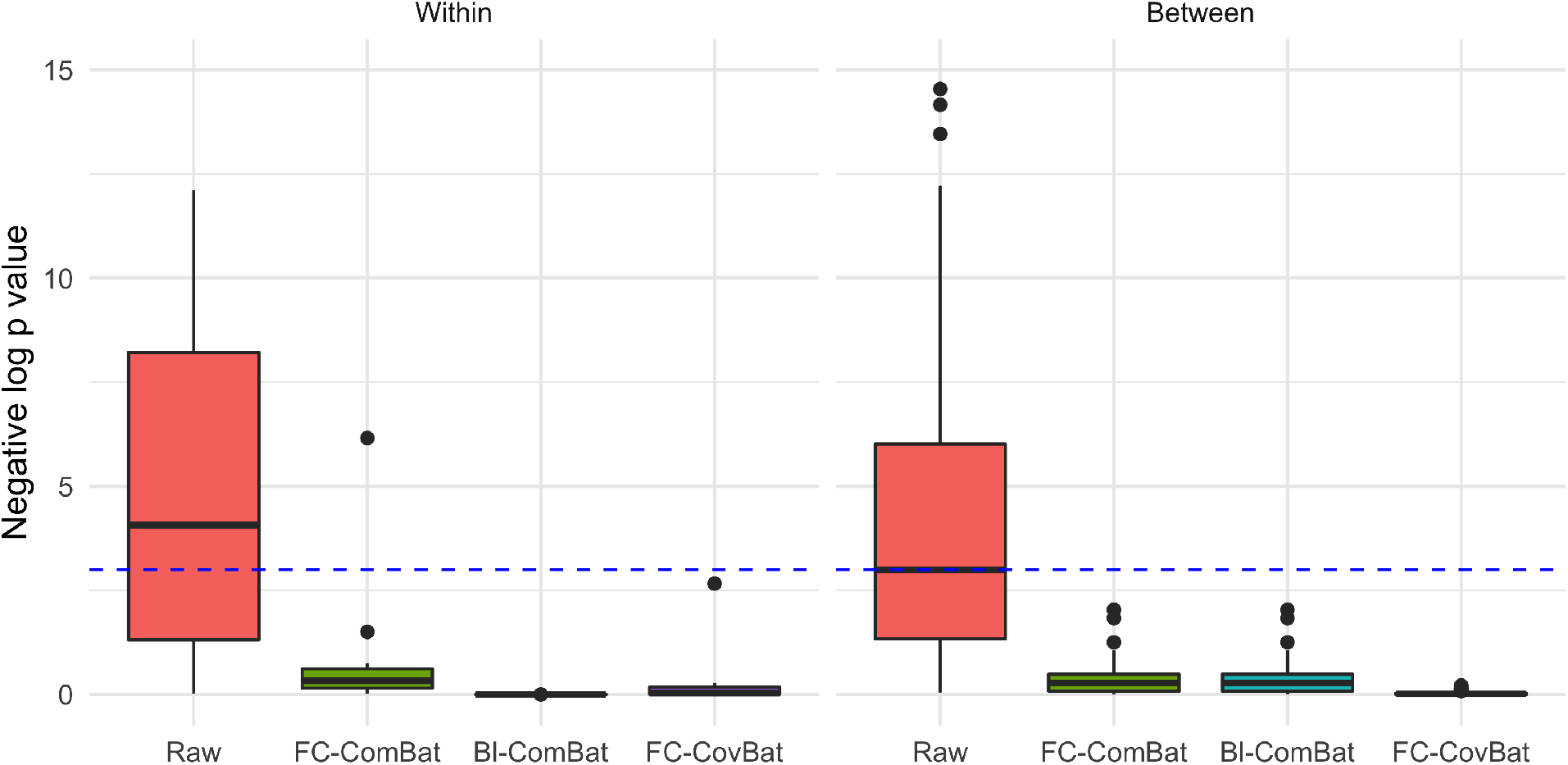
Scanner associations of within-network and between-network mean edge weights across subnetworks. Harmonization methods are compared in their reduction of scanner effects in within-network and between-network mean edge weights. Negative log *p*-values are shown for unharmonized and harmonized connectivities. The blue dotted line denotes the 0.05 significance threshold, where values higher than the line are considered significant.

### 3.3 Differences in scanner-average communities

In the unharmonized scanner-average communities, we find substantial differences across scanners (Fig. 5, mean pairwise ARI of 0.48). CARDIA-1 shows particularly notable differences, with the default mode network and sensory/somatomotor hand ROIs being distributed across more networks than other scanners. After any of the three harmonization methods though, these differences are substantially reduced, with FC-CovBat showing the greatest coherence across scanners (mean ARI of 0.74). We observe more severe scanner effects among WSBM-derived communities (mean ARI of 0.37; Supplementary Figure 1) with improved coherence after FC-ComBat (0.44), Bl-ComBat (0.47), and FC-CovBat (0.47).

**Figure 5:**
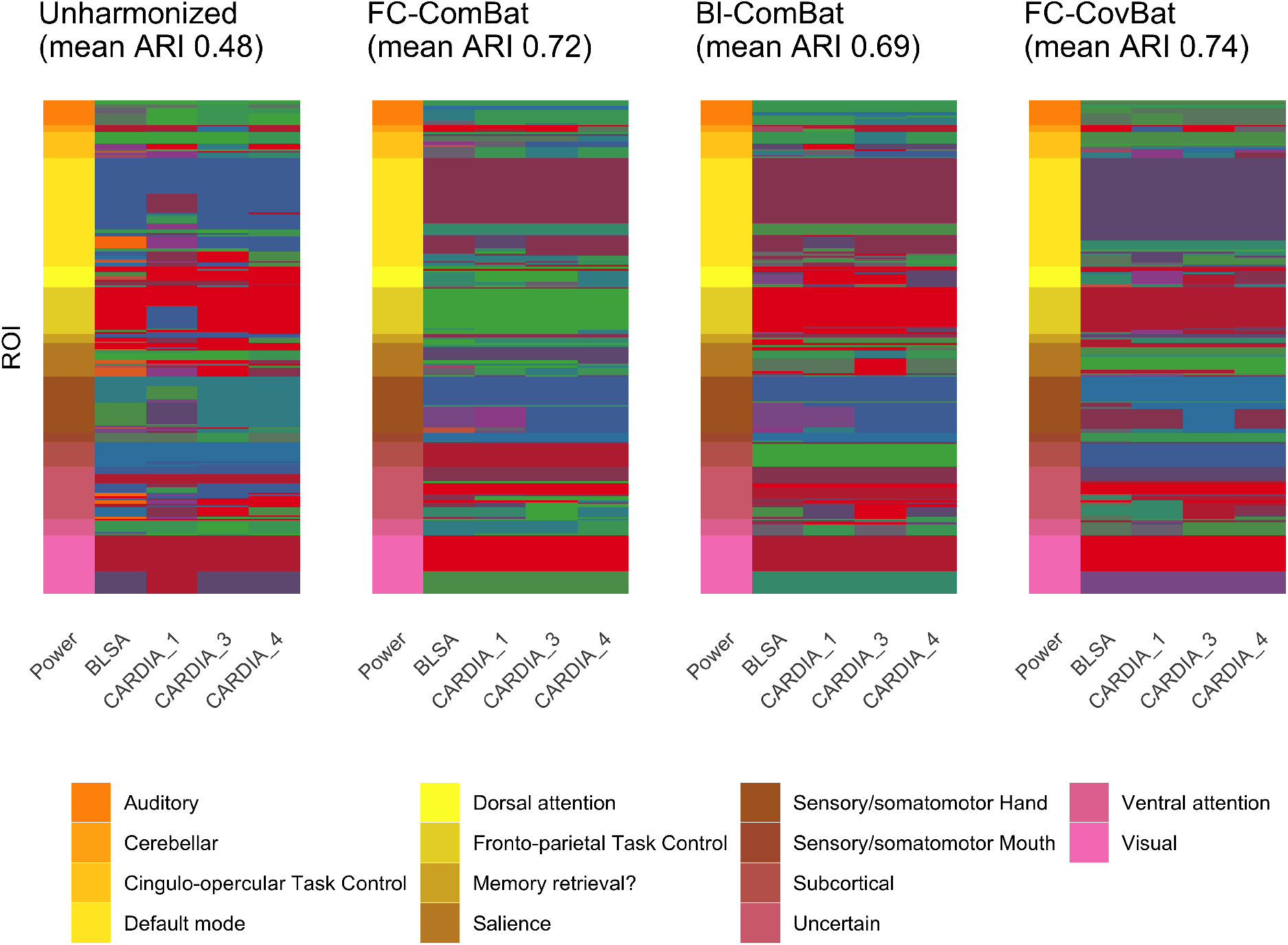
Signed Louvain communities in scanner-average networks before and after harmonization. Differences in scanner-specific communities are visualized across harmonization methods. The Louvain algorithm is used to derive communities from networks obtained by averaging the functional connectivity matrices across subjects acquired on each scanner. For numerical comparison, the adjusted Rand index is calculated between each partition and averaged to yield a measure of overall coherence.

### 3.4 Scanner effects in subject-specific communities

Communities at the subject level show notable scanner effects both within and across the two studies. PERMANOVA shows significant differences both for all scanners and limited to CARDIA scanners, with the latter being addressed by any of the harmonization methods (Fig. 6a). The remaining difference among all scanners after harmonization may be partially explained by demographic differences between the two studies (see Table 1). Distance-based regression via MDMR reveals that scanner influences subject-level communities, even while controlling for age and sex. While all harmonization methods reduce this scanner effect, only communities derived from FC-CovBat-adjusted data shows no significant effect of scanner (Fig. 6b).

**Figure 6:**
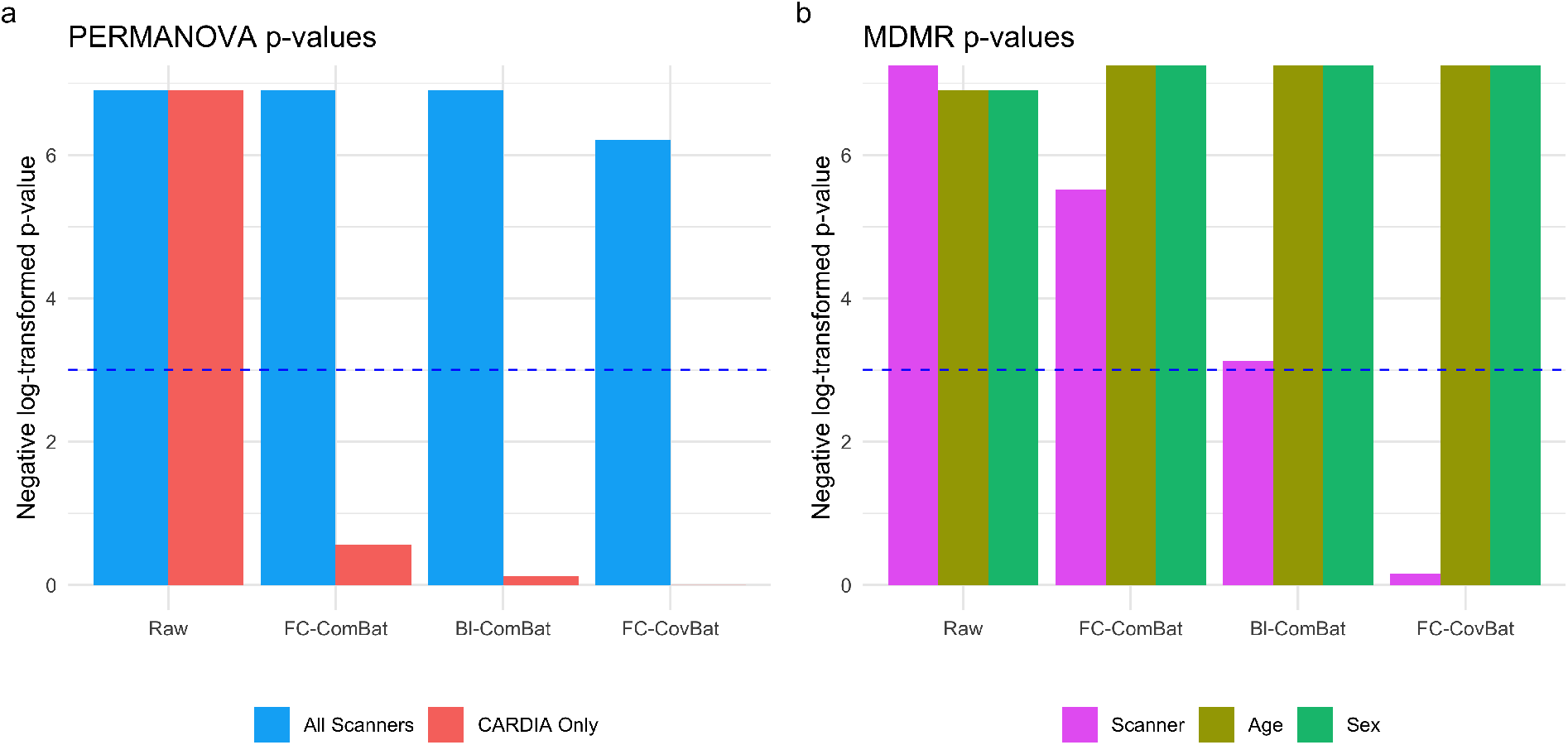
Assessment of scanner effects in subject-specific communities. Negative log *p*-values are displayed for permutational analysis of variance (PERMANOVA) in **a** and multivariate distance matrix regression (MDMR) in **b**. Both analyses compute distances as variation of information between subjects’ community partitions. Two separate PERMANOVAs test for group differences across all scanners and across CARDIA scanners only (CARDIA only). MDMR is performed once while including scanner, age, and sex in the model. The blue dotted line denotes the 0.05 significance threshold, where values higher than the line are considered significant.

### 3.5 Metrics of community structure

Across all network metrics considered, we find significant associations with scanner prior to harmonization (Table 2, **Supplementary Table 1**, and **Supplementary Table 2**)). Significant scanner associations with A. Mod., C. Mod., and CC are no longer present after any of the harmonization methods. The scanner effects on positive PC are significant for 7 of the subnetworks but no longer significant after any of the harmonization methods considered. Scanner effects in negative PC are similarly reduced (**Supplementary Table 2**). Significant age associations with both positive and negative PC are maintained after all of the harmonization methods considered (**Supplementary Table 1** and **Supplementary Table 2**). We also find a significant age association in A. Mod. which is maintained after harmonization, but no association was found with C. Mod. or CC (Table 2).

**Table 2:** Associations of network metrics with scanner and age before and after harmonization. Results from linear regression of metrics on scanner and age are displayed. *p*-values are shown for signed Louvain community modularity (C. Mod.), atlas modularity (A. Mod.), and clustering coefficient (CC)

|  | Scanner |  |  | Age |  |  |
| --- | --- | --- | --- | --- | --- | --- |
|  | C. Mod. | A. Mod. | CC | C. Mod. | A. Mod. | CC |
| Unharmonized | < 0.01 | < 0.01 | < 0.01 | 0.37 | < 0.01 | 0.54 |
| FC-ComBat | 0.76 | 0.86 | 0.13 | 0.44 | < 0.01 | 0.55 |
| Bl-ComBat | 0.69 | 1 | 0.1 | 0.32 | < 0.01 | 0.56 |
| FC-CovBat | 0.69 | 1 | 0.14 | 0.31 | < 0.01 | 0.52 |

## 4 Discussion

The increasing scale of rsfMRI studies merits particular consideration of scanner effects and their influence on studies of brain functional organization. We show considerable scanner effects throughout all of our evaluations, which include edge weights, community detection on multiple scales, and several metrics of network organization. We then demonstrate that several harmonization methods reduce scanner effects across our analyses, but Bl-CovBat further addresses scanner effects in edge weights and FC-CovBat is uniquely able to remove scanner effects in subject-specific communities.

Our discovery of scanner effects in community detection highlight additional concerns for future studies. For studies that derive networks from averaged connectivity matrices, our results suggest that community detection results could be influenced by the scanner used. When comparing community detection results across groups, researchers should be aware that differences could be driven by imbalance in subject demographics across scanners. For studies focused more on subject-level analyses, our findings suggest that each subject’s community detection results may be partially driven by scanner, which again may confound study results.

Our analyses of network metrics replicate previous findings while finding novel scanner effects. We again identify major scanner effects in within and between-network connectivity values, which are largely addressed through harmonization (Yu et al., 2018). We find age associations in within-subnetwork mean edge weights and average positive participation coefficient, both consistent with previous studies (Yu et al., 2018; Varangis et al., 2019). We identify scanner effects in metrics of network organization that have previously not been considered, including maximum modularity from the Louvain method, modularity with respect to the atlas, clustering coefficient, and participation coefficient. Scanner effects in maximum modularity achieved through the Louvain method suggest community detection algorithms may vary in performance depending on the scanners used. Furthermore, scanner differences in modularity could suggest that the fit of functional atlases may depend on scanner properties of studies using these atlases.

The harmonization methods generally perform well on all evaluations, but Bl-ComBat and FC-CovBat outperform FC-ComBat in certain scenarios. FC-ComBat performs well on harmonizing the subject-average networks since it removes scanner effects in the mean of edge weights. However, FC-CovBat is necessary to fully address effects in the subject-specific networks, where scanner effects in groups of edges may drive differences in each individual’s community structures. Compared to the other methods, Bl-ComBat performs best for removing within-network scanner effects owing to its additional ComBat steps within each specified block. However, Bl-ComBat may not be applicable in some studies since blocks must be specified a priori, which is not feasible if the ROIs are not defined based on previous network analyses. FC-CovBat does not require any prior information and provides the greatest removal of scanner effects in between-network connectivities and subject-specific community structures.

There are several limitations with the current study that worth future investigation and extensions. Our proposed harmonization methods build on the ComBat framework that treats edges as individual features and may not preserve the positive semidefinite property of the functional connectivity matrices. While lacking this positive semidefinite property does not prevent construction of networks from the matrices, several analyses are no longer possible after application of our harmonization methods. For instance, both covariance regression techniques (Chiu et al., 1996; Hoff & Niu, 2012; Zhao et al., 2019; Zou et al., 2017) and analyses on the Riemannian space of positive semidefinite matrices (You & Park, 2021) cannot be applied after our harmonization procedures. To enable those analyses on harmonized data, future studies could investigate removal of scanner effects through spectral models (Flury, 1984; Zhao et al., 2019) or transformations (Chiu et al., 1996), which would both preserve the positive semidefinite property.

Since we employ network analysis methods that are prevalent throughout the literature and relevant to our aims, our evaluations do not reflect some recent advances in network methodologies. We define ROIs and subnetworks using the Power atlas (Power et al., 2011) but more recent atlases might provide different parcellations and could capture different information from the functional imaging (Eickhoff et al., 2018). In examining population-average communities, we construct our weighted networks using the arithmetic mean of functional connectivity matrices after Fisher-transformation of the entries. However, a previous study has shown that this averaged functional connectivity matrix may not retain properties of the individual matrices and proposes exponential random graph model to yield a group-based network that better preserves individual properties (Simpson et al., 2012). In applying the Louvain method and the WSBM, we choose algorithm parameters to match the number of subnetworks in the atlas. However, methods exist to find parameters that optimally fit the networks, including CHAMP (Convex Hull of Admissible Modularity Partitions; Weir et al., 2017) for modularity maximization algorithms and Bayes factors for WSBMs (Aicher et al., 2015). Future studies could investigate scanner effects using these data-adaptive network analysis techniques.

In our study, we examine a single study design for deriving networks and communities from rsfMRI observations and future studies could test different study designs. Instead of using correlation values to construct edge weights, studies could examine how partial correlations could influence downstream scanner effects (Marrelec et al., 2006). Estimating sparse networks (Friedman et al., 2008) could also aid in reducing scanner effects, but this direction has yet to be explored. While we considered communities obtained via the Louvain algorithm and WSBMs, many other algorithms could also be considered (Leskovec et al., 2010; Traag et al., 2019). The choice of atlas has previously been demonstrated to influence scanner effects in network metrics (Yu et al., 2018), but this could be extended to examine how scanner effects in communities could also be affected. While different network construction methods may have varying dependence on scanner, our study suggests that studies of brain functional organization should consider scanner as a potential confound and follow-up studies could further reveal the breadth of this issue.

## Data and code availability statement

The data that support the findings of this study are not publicly available due to restrictions imposed by the administering institution and privacy of the participants. The authors will share them by request from any qualified investigator after completion of a data sharing agreement.

The harmonization methods used in the paper are implemented in the FCHarmony package available at https://github.com/andy1764/FCHarmony. Network analyses are performed using the Brain Connectivity Toolbox (BCT) version 2019-03-03 available on their website at https://sites.google.com/site/bctnet/.

## CRediT authorship contribution statement

R.T.S. and H.S. contributed equally to this work. **Andrew A. Chen:** Conceptualization, Formal analysis, Methodology, Software, Investigation, Writing - Original Draft. **Dhivya Srinivasan:** Data Curation, Writing - Review & Editing. **Raymond Pomponio:** Data Curation. **Yong Fan:** Data Curation, Supervision. **Ilya M. Nasrallah:** Data Curation, Supervision, Project Administration. **Susan M. Resnick:** Data Curation, Supervision, Project Administration. **Lori L. Beason-Held:** Data Curation, Supervision, Project Administration. **Christos Davatizikos:** Data Curation, Supervision, Project Administration. **Theodore D. Satterthwaite:** Conceptualization, Writing - Review Editing. **Dani S. Bassett:** Conceptualization, Writing - Review Editing. **Russell T. Shinohara:** Supervision, Conceptualization, Methodology, Writing - Review Editing. **Haochang Shou:** Supervision, Conceptualization, Methodology, Writing - Review Editing.

## Acknowledgements

This work was supported by the National Institute of Neurological Disorders and Stroke (grant numbers R01 NS085211 and R01 NS060910), the National Multiple Sclerosis Society (RG-1707-28586) and a seed grant from the University of Pennsylvania Center for Biomedical Image Computing and Analytics (CBICA). The content is solely the responsibility of the authors and does not necessarily represent the official views of the funding agencies.

The Baltimore Longitudinal Study of Aging is supported by the Intramural Research Program of the National Institute on Aging, NIH.

