## Supplementary materials for "Harmonizing Functional Connectivity Reduces Scanner Effects in Community Detection"

[a] *Penn Statistics in Imaging and Visualization Center, Department of Biostatistics, Epidemiology, and Informatics, University of Pennsylvania, Philadelphia, PA 19104, USA*
[b] *Center for Biomedical Image Computing and Analytics, University of Pennsylvania, Philadelphia, PA 19104, USA*
[c] *Department of Radiology, University of Pennsylvania, Philadelphia, PA 19104, USA*
[d] *Department of Biostatistics, Colorado School of Public Health, Aurora, CO 80045, USA*
[e] *Laboratory of Behavioral Neuroscience, National Institute on Aging, Baltimore, MD 21224, USA*
[f] *Penn Lifespan Informatics & Neuroimaging Center, Department of Psychiatry, University of Pennsylvania, Philadelphia, PA 19104, USA*
[g] *Department of Psychiatry, University of Pennsylvania, Philadelphia, PA 19104, USA*
[h] *Department of Bioengineering, School of Engineering  Applied Science, University of Pennsylvania, Philadelphia, PA 19104, USA*
[i] *Department of Electrical  Systems Engineering, School of Engineering  Applied Science, University of Pennsylvania, Philadelphia, PA 19104, USA*
[j] *Department of Physics  Astronomy, College of Arts  Sciences, University of Pennsylvania, Philadelphia, PA 19104, USA*
[k] *Department of Nuerology, Perelman School of Medicine, University of Pennsylvania, Philadelphia, PA 19104, USA*
[l] *Department of Psychiatry, Perelman School of Medicine, University of Pennsylvania, Philadelphia, PA 19104, USA*
[m] *Santa Fe Institute, 1399 Hyde Park Rd, Santa Fe, NM 87501, USA*
[1] *Equal contribution*

***Correspondence: Andrew A. Chen**


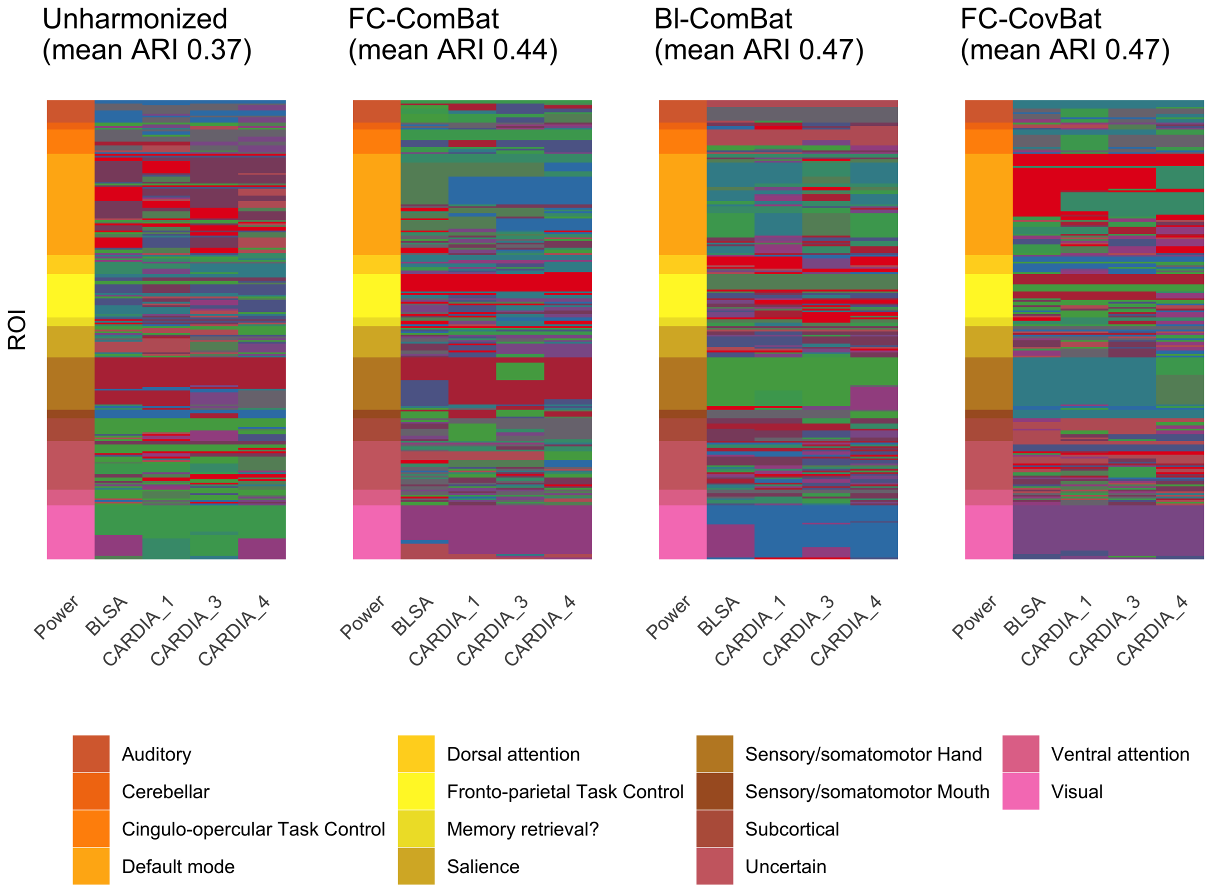


Supplementary Figure 1: **Weighted stochastic block model communities in scanner-average networks before and after harmonization.** A weighted stochastic block model (WSBM) is used to derive communities from networks obtained by averaging the functional connectivity matrices across subjects acquired on each scanner. The WSBM is fit to identify 14 communities. For numerical comparison, adjusted Rand index is calculated between each pair of scanners and averaged to yield a measure of overall coherence.

| | Scanner | | | | Age | | | |
|---|---|---|---|---|---|---|---|---|
| | Unharmonized | FC-ComBat | Bl-ComBat | FC-CovBat | Unharmonized | FC-ComBat | Bl-ComBat | FC-CovBat |
| Auditory | 0.03 | 0.33 | 0.46 | 0.34 | 0.78 | 0.87 | 0.88 | 0.86 |
| Cerebellar | 0.1 | 0.93 | 0.75 | 0.93 | 0.64 | 0.65 | 0.65 | 0.63 |
| Cingulo-opercular Task Control | 0.05 | 0.19 | 0.42 | 0.17 | 0.76 | 0.34 | 0.37 | 0.31 |
| Default mode | $< 0.01$ | 0.49 | 0.99 | 0.86 | $< 0.01$ | $< 0.01$ | $< 0.01$ | $< 0.01$ |
| Dorsal attention | 0.65 | 0.94 | 0.75 | 0.94 | 0.64 | 0.93 | 0.93 | 0.92 |
| Fronto-parietal Task Control | 0.69 | 0.91 | 0.84 | 0.87 | 0.01 | $< 0.01$ | $< 0.01$ | $< 0.01$ |
| Memory retrieval | $< 0.01$ | 0.27 | 0.94 | 0.28 | $< 0.01$ | $< 0.01$ | $< 0.01$ | $< 0.01$ |
| Salience | 0.04 | 0.28 | 0.25 | 0.26 | 1 | 0.84 | 0.87 | 0.81 |
| Sensory/somatomotor Hand | 0.13 | 0.53 | 0.8 | 0.63 | 0.01 | 0.01 | 0.01 | 0.01 |
| Sensory/somatomotor Mouth | 0.39 | 0.74 | 0.77 | 0.76 | 0.02 | 0.02 | 0.02 | 0.02 |
| Subcortical | 0.01 | 0.28 | 0.74 | 0.26 | 0.8 | 0.96 | 0.98 | 0.98 |
| Uncertain | 0.36 | 0.59 | 0.8 | 0.59 | 0.01 | 0.01 | 0.01 | 0.01 |
| Ventral attention | 0.01 | 0.22 | 0.28 | 0.21 | 0.56 | 0.58 | 0.58 | 0.6 |
| Visual | 0.02 | 0.26 | 0.88 | 0.76 | 0.62 | 0.6 | 0.59 | 0.58 |

Supplementary Table 1: **Associations of positive participation coefficient with age and site.** Positive participation coefficient (PC) is calculated for each node defined by the Power atlas and averaged within Power atlas subnetworks. These averaged positive PC values are regressed on site and age and the corresponding $p$-values for each regressor are displayed. $p$-values significant at the 0.05 level are highlighted in blue.

| | Scanner | | | | Age | | | |
|---|---|---|---|---|---|---|---|---|
| | Unharmonized | FC-ComBat | Bl-ComBat | FC-CovBat | Unharmonized | FC-ComBat | Bl-ComBat | FC-CovBat |
| Auditory | 0.07 | 0.37 | 0.82 | 0.34 | $< 0.01$ | $< 0.01$ | $< 0.01$ | $< 0.01$ |
| Cerebellar | $< 0.01$ | 0.03 | 0.28 | 0.03 | $< 0.01$ | $< 0.01$ | $< 0.01$ | $< 0.01$ |
| Cingulo-opercular Task Control | $< 0.01$ | 0.26 | 0.77 | 0.25 | $< 0.01$ | $< 0.01$ | $< 0.01$ | $< 0.01$ |
| Default mode | $< 0.01$ | 0.47 | 0.58 | 0.49 | 0.18 | 0.19 | 0.09 | 0.21 |
| Dorsal attention | $< 0.01$ | 0.14 | 0.67 | 0.15 | $< 0.01$ | 0.01 | 0.01 | 0.01 |
| Fronto-parietal Task Control | 0.04 | 0.2 | 0.46 | 0.2 | 0.01 | $< 0.01$ | $< 0.01$ | 0.01 |
| Memory retrieval | 0.76 | 0.56 | 0.62 | 0.57 | 0.93 | 0.83 | 0.83 | 0.78 |
| Salience | 0.01 | 0.16 | 0.53 | 0.15 | $< 0.01$ | $< 0.01$ | $< 0.01$ | $< 0.01$ |
| Sensory/somatomotor Hand | 0.27 | 0.6 | 0.63 | 0.65 | 0.23 | 0.24 | 0.24 | 0.26 |
| Sensory/somatomotor Mouth | $< 0.01$ | 0.1 | 0.26 | 0.1 | $< 0.01$ | $< 0.01$ | $< 0.01$ | $< 0.01$ |
| Subcortical | $< 0.01$ | 0.08 | 0.11 | 0.14 | $< 0.01$ | $< 0.01$ | $< 0.01$ | $< 0.01$ |
| Uncertain | 0.16 | 0.14 | 0.25 | 0.16 | 0.07 | 0.08 | 0.07 | 0.07 |
| Ventral attention | 0.02 | 0.22 | 0.47 | 0.21 | $< 0.01$ | $< 0.01$ | $< 0.01$ | $< 0.01$ |
| Visual | $< 0.01$ | 0.1 | 0.68 | 0.09 | $< 0.01$ | $< 0.01$ | $< 0.01$ | $< 0.01$ |

Supplementary Table 2: **Associations of negative participation coefficient with age and site.** Negative participation coefficient (PC) is calculated for each node defined by the Power atlas and averaged within Power atlas subnetworks. These averaged negative PC values are regressed on site and age and the corresponding $p$-values for each regressor are displayed. $p$-values significant at the 0.05 level are highlighted in blue.
